# Formation, stability, and competition in dendritic proto-spines

**DOI:** 10.64898/2026.01.09.697842

**Authors:** V. V. Pulikkottil, U.S. Bhalla

**Affiliations:** National Center for Biological Sciences, Tata Institute of Fundamental Research, Bangalore, Karnataka, India

**Author notes:** VVP and USB designed the project, wrote the paper and obtained funds. VVP conducted the simulations, USB supervised the project. The authors declare that they have no competing interests.

**Keywords:** synaptic spines, IBAR proteins, spine spacing, Neurodevelopmental disorders

## Abstract

Over 80 % of excitatory synapses in the mammalian brain terminate on dendritic spines, which support key functions of memory storage and computation. Despite the neurodevelopmental and computational implications of spine formation, little is known about the interplay of physical and chemical signaling mechanisms that govern spine initiation. We developed a physics-based model for the formation, stability, and placement of membrane protrusions (proto-spines) triggered by membrane-bending proteins such as IRSp53, and activated by neurotransmitter inputs. We identify two distinct proto-spine energy manifolds depending on the concentration of activated IRSp53, causing shallow and sharp geometries respectively. The sharp proto-spines exhibit properties of wave-pinning models, including stimulus-local formation, positional stability, and merging, but unlike the wave-pinning model they also exhibit competition. We linked the physics-based model to a mass-action chemistry model of neurotransmitter-driven signaling pathway activation of IRSp53 to simulate proto-spine formation in response to neurotransmitter stimulus patterns. Multiple proto-spines compete for IRSp53, leading to inter-spine spacing of 1 - 3 *µm*, similar to experiments. Correlated and spatially clustered synaptic input enhances proto-spine lifetime and density compared to non-correlated stimuli. Finally, we tested the effects of autism spectrum disorder-related mutations in spine-formation signaling, leading to altered RhoGEF activity. We found that elevated RhoGEF activity increased proto-spine density and reduced their lifetime, consistent with ASD phenotypes. Overall, our analysis shows that the interplay between membrane bending mechanics and reaction-diffusion chemistry creates rich proto-spine dynamics and concisely captures key features of early spine formation.

**Significance Statement:** Dendritic spines mediate most of the excitatory neurotransmission in the brain, and alterations in their shape and distribution on the dendrite are correlated with neuro-developmental and neuro-degenerative conditions. We model membrane mechanics and chemical signaling to show that spine precursors arise as self-stabilizing membrane protrusions due to the binding of membrane-bending proteins. Spine precursors act as nucleation sites for spine-building proteins, and hence their stability, placement, and signaling impacts late-stage spine maturation. These proto-spines compete with each other for bending proteins, and we connect this interplay with signaling chemistry that triggers new spines immediately below sites of transmitter release. We find that aberrant signaling in neurological disease alters proto-spine competition, life-time and placement.

---

The dendritic spine is a key unit of dendritic computation in pyramidal cells of the cortex and hippocampus, in cerebellar Purkinje neurons, and numerous other neuronal types (1–3). It mediates neuronal information processing by compartmentalizing signaling and plasticity, thus controlling the information flow to the dendrite. Spines in clusters enhance cellular responses by amplifying subthreshold electrical signals, and share signaling molecules leading to heterosynaptic plasticity (4–8).

Spines are dynamic. They form in response to presynaptic neurotransmitter release or glutamate uncaging (9), and spine turnover is implicated in synaptic plasticity (10). Spine density changes occur in response to stress (11), sleep (8), and enriched environments (12, 13). Neurotransmitters trigger structural plasticity and diffusional cross-talk through Rac1 and Cdc42 between spines spaced within 5 *µm* (14, 15). Glutamate uncaging stimuli of 40 pulses at 0.5 Hz have been shown to cause de novo spine formation (9) in an NMDAR-dependent manner, leading to regulation of spine plasticity and density changes in CNS (16). The signaling substrate for this includes glutamate-dependent pathways, which connect NMDAR to structural plasticity proteins Rac1, Cdc42, and RhoA through CaMKII (15, 17–19), facilitating spine formation and diffusional cross-talk among neighbouring spines. These plasticity proteins and their modulators, such as Tiam1 (20) and Trio (21), have been implicated in altered spine structure and distribution in neurodevelopmental and neurodegenerative disorders (22). Despite the overlap in signaling pathways involved in spine formation and synaptic plasticity, most spine formation models focus on actin-induced structural mechanisms (23–27), whereas numerous synaptic plasticity models focus on chemical signaling (28–31).

Spine formation is a multi-stage process, beginning with synaptic activity-induced membrane deformation (32, 33). This membrane curvature is mediated by proteins with a curved I-BAR domain, such as IRSp53 and MIM (34–36). IRSp53 can bind to flat giant unilamellar vesicle (GUV) membranes, cluster with actin regulators like vasodilator-stimulated phosphoprotein (VASP) (37), and cause membrane protrusions (38). Several lines of evidence implicate IRSp53 in early spine formation. It colocalizes with proteins related to actin dynamics, PSD-95 (39, 40), and SHANK (39, 41). Rac-1 and Cdc42, which are known modulators of spinogenesis (28, 42) activate IRSp53 in response to extracellular stimuli. The knockdown of IRSp53 causes anomalies in the morphology and distribution of spines (43).

Cell membranes are supported by a cytoskeletal network. In axons this is provided by a membrane associated periodic skeleton (MPS) spaced at 190 nm (44), and some of these proteins are also present in dendrites (45). MPS is remodeled by calcium signaling (44), involving Rac-1 and Cdc42, which also modulate IRSp53 activity. Actin cytoskeletal rearrangement can also happen as a result of the combined effect of I-BAR proteins and Rac-1 (46). Thus the dynamics of dendritic membrane shape are liable to be affected by IRSp53 signaling in coordination with actin remodeling.

Spine formation theories typically begin with actin dynamics (23, 47), and incorporate few details of the steps above. However, several experiments link membrane protrusions and spine precursors, such as filopodia (48), to later actin mechanisms (49). Spine precursors accumulate molecules that bind actin, actin elongating and branching proteins, and actin severing proteins to form a mature spine (33, 50). The shape of a mature spine is integral to its function due to its role in isolating postsynaptic signaling and bringing together proteins that respond to neurotransmitter release (51, 52).

Several factors contribute to the energy budget of a membrane protrusion. Protein-induced membrane bending incurs a mechanical energy cost (53). This arises from the mismatch between intrinsic protein curvature and induced membrane curvature, which is mathematically quantified using Helfrich energy (54). Protein-protein interactions on the membrane contribute to competing entropy and aggregation terms (55, 56). The concentration gradient of IRSp53 between the dendritic cytosol and membrane of the proto-spine drives molecular exchange between the cytosol and the membrane(57, 58). The curvature sensing of IRSp53 also recruits them to preferred membrane curvatures (59). A range of models have looked at membrane protrusion (55, 60, 61), but to our knowledge, none have integrated all these contributions. The exchange term, in particular, is crucial to simulate neurotransmitter signaling-driven membrane protrusion.

Here, we combined a physics-based model incorporating protein-membrane interaction energies with a model of the chemical processes downstream of spatiotemporal stimulus patterns to collectively examine the dynamics of proto-spine initiation and distribution.

## Results

We modeled mechano-chemical processes of spine formation in the following stages: First, we developed a mechanical deformation model to characterize the energy landscape for IRSp53-induced proto-spine shapes. Second, we showed the similarities and differences between our model and existing biological pattern formation models such as wave-pinning and Turing patterns. Third, we simulated competition between proto-spines. Fourth, we coupled the mechanical model to chemical signaling, increasing biological detail in our model to describe phenomena arising from multi-scale cellular physiology. Finally, we showed how stimulus patterning and chemical pathway mutations affect spine density, size, and lifetimes in healthy and disease conditions.

### Energy minimization predicts biologically reasonable pro-to-spine size

Our first-pass model incorporated energy terms for membrane deformation in a quasi-two-dimensional analysis along the axial and radial dimensions of a dendrite. We included contributions from membrane-IRSp53 curvature mismatch, mechanical membrane bending, entropy, and IRSp53 aggregation energy (Methods). We treated proto-spines as spherical dome-shaped protrusions. The bending protein IRSp53 was confined uniformly to this dome. The dome merged into the locally flat dendritic membrane with a ring of negatively curved membrane (Fig. 1 A, B). We minimized energy with respect to the number of molecules in the membrane (*ϕ*_*tot*_). Unexpectedly, we found two minimal energy configurations (Fig. 1 E and supplementary movie S1 A, B): one corresponding to a sharp protrusion and another for a shallow protrusion (Fig. 1 G). The shallow and sharp branches are characterized by two distinct ranges of *ϕ*, i.e., *<*=0.1 and *>*0.3. Raising the IRSp53 concentration from 0.1 *µM* to 0.6 *µM* mostly eliminated the shallow energy branch (Fig. 1: F, supplementary movie S2 A, B). The diameter of the sharp protrusion (0.5 *µm* - 1.0 *µm*) resembles that of a filopodium, and the height (100*nm*) is larger than dendritic surface irregularities. Both of these attributes make the sharp protrusion a biologically plausible spine precursor.

**Fig. 1.**
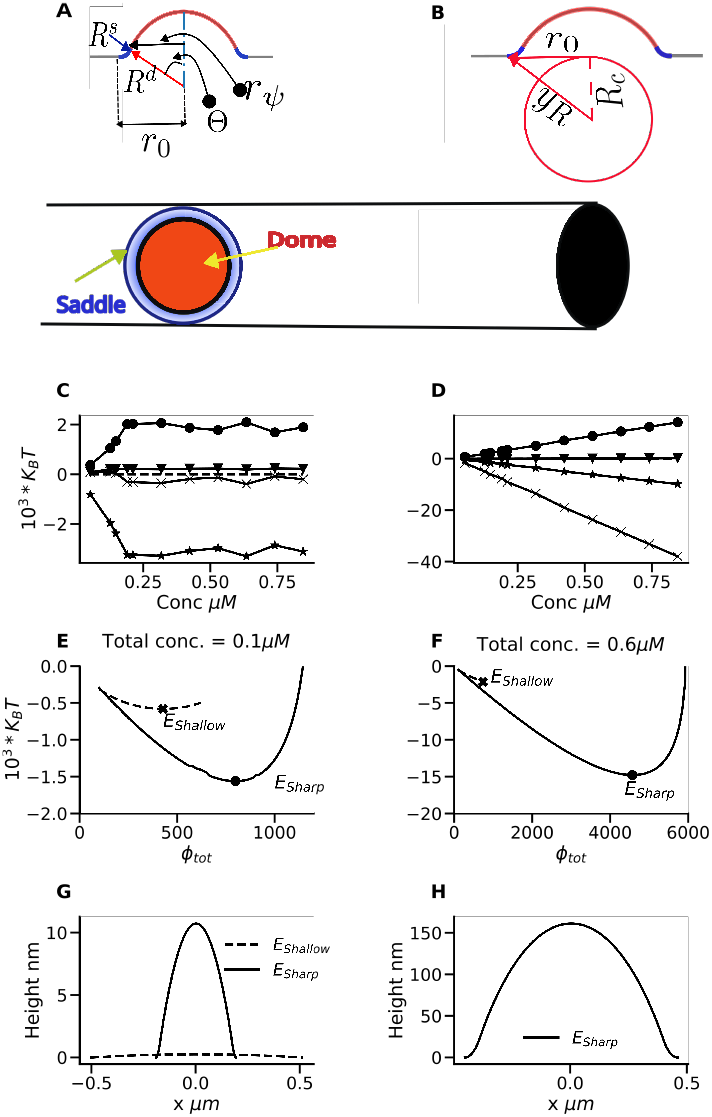
Proto-spines are shaped by distinct energy branches. A) Model layout with axisymmetric membrane protrusion. Top: *R*^*d*^ and *R*^*s*^ are dome and saddle radii, respectively. Θ is the angle between the line that connects the dome-saddle inflection point with the dome’s center and the vertical axis. *r*_*ψ*_ and *r*_0_ are derived from the *R*^*d*^, *R*^*s*^ and Θ. B) To keep protrusion width below the dendritic diameter, we introduce an energy cost (7) as a function of *sagging length yR*, C) and D) Contributions of each of the energy terms, as a function of IRSp53 concentration. (C)shows the energy contributions for the shallow low *ϕ* branch and (D) shows the energy contributions for the sharp high *ϕ* branch. The symbols used are as follows: “x” - Aggregation, “*” - Entropy, “o” - Mismatch, “v” - Constraint energies. E and F) Summed energy profiles for shallow and sharp energy branches, in two concentration ranges. G) Shallow and sharp shapes corresponding to 0.1 *µM* concentration. H) Sharp shape at 0.6 *µM* concentration.

#### Protrusion stability criteria match those from wave-pinning models

Turing models (62, 63) and wave-pinning models (64, 65) are classical spatiotemporal patterning models with local peaks. We stipulated that our model should have three key qualitative features: The position of the peaks should match the stimulus points; additional peaks should not alter the position of extant ones; and peaks should not spontaneously drift. The wave-pinning meets these criteria but not Turing models (Fig. 2 A, B). The wave-pinning criterion for stability is that the wave speed (speed of propagation of edges of the peak) should be zero (64) (Fig. 2 C). Furthermore, in the wave-pinning model, intersecting wave-pinning peaks merge (Fig. 2 D). Similar to wave-pinning, the wave speed for our model depends on the number of molecules in the protrusion *ϕ*_*tot*_ (Fig. 2 E). Absolute total energy (*E*_*sharp*_) as plotted in Fig. 1 E and F matches well with the ones obtained from wave speed analysis for a range of IRSp53 concentrations (Fig. 2 F). The resultant proto-spine widths (Fig. 2 H) are similar to the spine widths in the literature (66, 67). Bulk IRSp53 concentration controls *ϕ*_*tot*_ indirectly through the energy minimization conditions (Methods: shape optimization). We performed a numerical simulation to model diffusion of molecules in the bulk which are also recruited to the membrane. We ran this calculation till steady-state, which should match the energy minimum. We found a roughly linear relation between energy and concentration, and the simulated output agreed both with the analytic absolute energy minimum *E*_*sharp*_ (Fig. 1 E, F), and with the solution from the analytic criterion for wave-pinning stability (Fig. 2 F). The *ϕ*_*tot*_ dependence on IRSp53 concentration was sensitive to the protein aggregation coefficient *K*_*agg*_ (Fig. 2 G), and so was the size of the resultant proto-spine (Fig. 2 H). Thus in this section we had established the similarity between wave-pinning patterning and our model, and obtained biologically plausible size ranges for the resultant proto-spines.

**Fig. 2.**
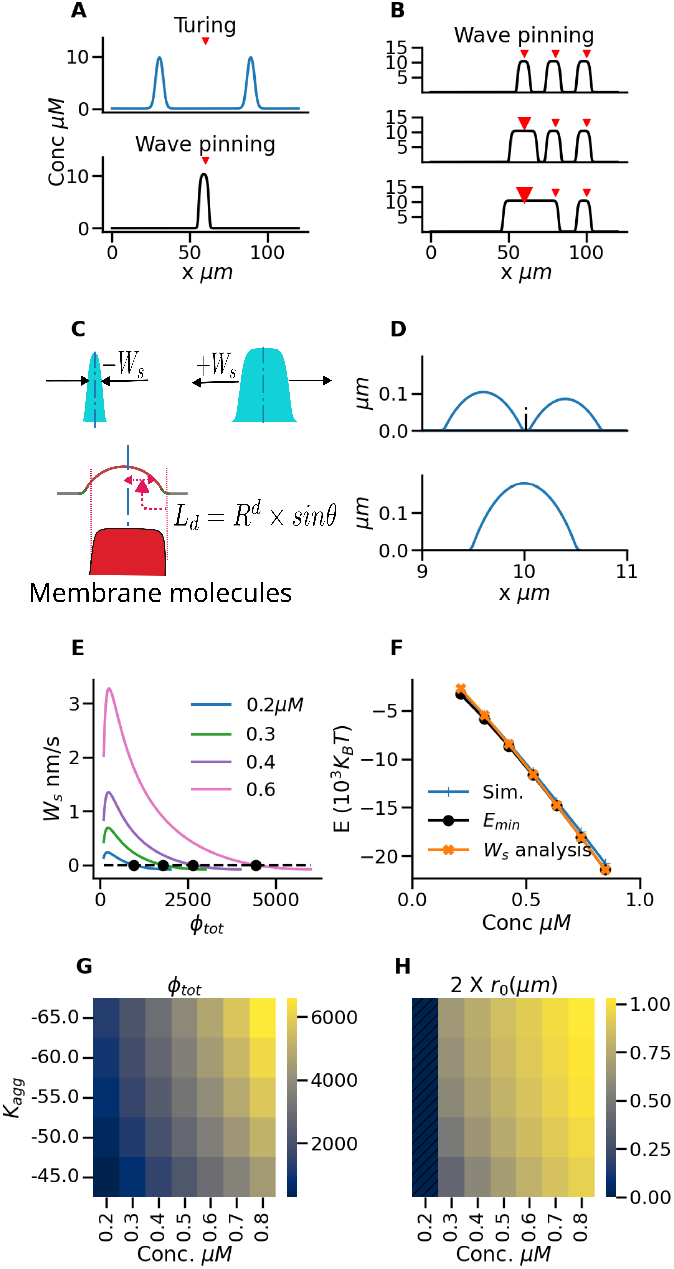
Spine-protrusion model fits wave-pinning but not Turing model properties. A) Turing pattern peaks (top) do not necessarily align to stimulus location, but wave-pinning model peaks (bottom) do align. In A) and B) red arrows represent the stimulus locations. B) Wave-pinning produces multiple local peaks under stimuli (top) which get larger with stronger stimuli (middle), and overlapping peaks merge (lower). C) The membrane front propagates with a positive or negative velocity depending on the number of molecules (*ϕ*_*tot*_) attached to the membrane. D) In the spine-protrusion model, we detect proto-spine intersection programmatically and then construct a single proto-spine with the summed number of IRSp52 molecules. E) Wave speed in spine protrusion model is a function of the total number of molecules in the membrane, i.e., *ϕ*_*tot*_. F) Comparison of three methods to estimate energy minimum as a function of concentration of IRSp53: simulation, wave-speed analysis, and the absolute energy minimum (*E*_*sharp*_). G) Dependence of *ϕ*_*tot*_ on IRSp53 aggregation within the membrane (*K*_*agg*_) and the bulk IRSp53 concentration. H) Spine size (in *µm*) varies with the aggregation coefficient and bulk IRSp53 concentration.

### Resource Competition leads to a minimum spacing requirement between proto-spines

Adjacent spines are coupled by diffusion of IRSp53. This potentially leads to cooperation to form clusters (6, 7, 10), or to competition and spine loss (10). The diffusion length-scale of 5 - 10*µm* (29) is similar to the spacing between co-active spines in a cluster (6 *µm* (68)). To explore this with our model, we initiated four proto-spines, symmetrically placed about the center of the 20 *µm* long dendrite. We varied inter-spine spacing from 1.5 *µm* down to 0.55 *µm*. We found there was an intermediate range of spine spacing and IRSp53 concentration over which all four spines persisted (Fig. 3 A). As expected, under resource-limited conditions (low concentration of IRSp53), competition led to elimination of spines. There were two opposing effects when spines were crowded, i.e., for lower inter-spine spacings. At near threshold IRSp53 levels (*<* 0.48 *µM*) spines competed and fewer survived. Conversely, in resource-rich conditions, the proto-spines grew till they merged with each other (Fig. 3 A). When multiple proto-spines did co-exist, they were much more shallow than cases where spine elimination had occurred (Fig. 3 B, C).

**Fig. 3.**
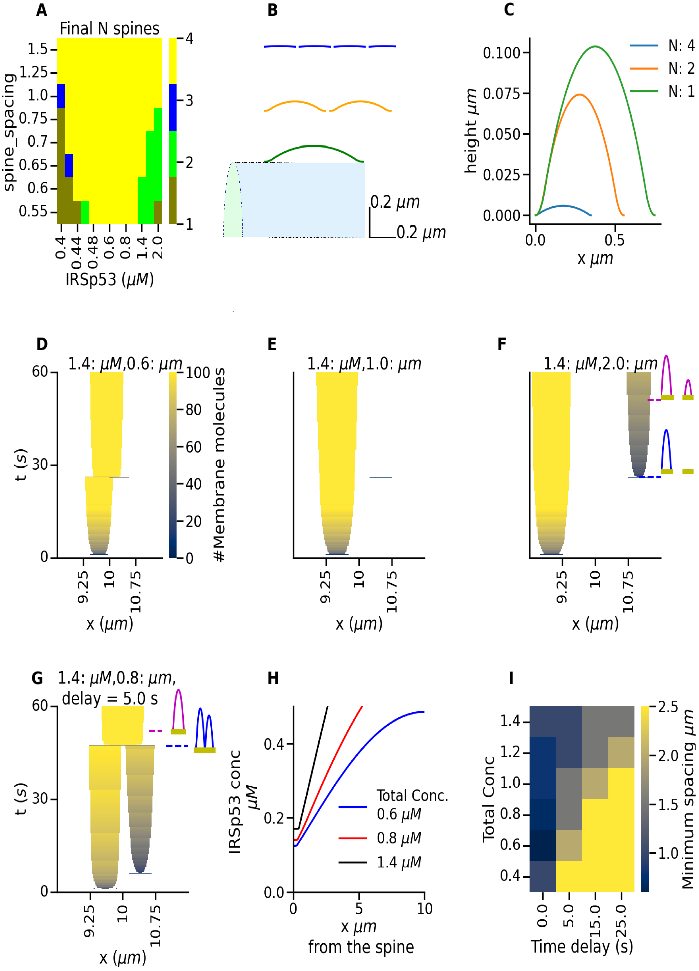
Proto-spine spacing is determined by IRSp53 availability and initiation time. A) Map of domains of spine survival following initiation of four equally-spaced spines. At closer spacing, both spine competition and spine merger contribute to spine loss, depending on IRSp53 levels. B,C) Proto-spines become shallower when more are formed at the same concentration of IRSp53. B) Scale with respect to dendrite. C) Proto-spine shapes overlaid. D-G) Growth profiles of spine interactions. When a second proto-spine is initiated with time delay = 25 seconds, it D) merges immediately if spacing *<* the proto-spine diameter, E) loses to competition for small spacings, F) survives for large spacing. In G, the second proto-spine coexists briefly before merging when it is created with a shorter delay of 5 s after the first. H) Dendrite concentration of IRSp53 for distance ‘x’ away from the center of the first proto-spine at time t = 60s after spine initiation. I) Minimum spine spacing for different initiation delays and concentrations.

Do pre-existing spines have a competitive advantage? We simulated formation of two proto-spines 25 seconds apart, with a range of spacings. As expected, the larger, pre-existing proto-spine was more stable and consumed the new spines if they were too close (Fig. 3 E). Interestingly, at high IRSp53 concentrations, it was possible for a new spine to grow briefly, but then merge with an existing spine to form a combined one at an intermediate position (Fig. 3 G). To our knowledge, merging does not happen between mature spines, hence this could be a unique feature of proto-spine dynamics. There was a steep fall-off of available IRSp53 in the dendrite beneath the existing proto-spine (Fig. 3 H). This creates a zone in which there is insufficient IRSp53 (*<* 0.4 *µM*) to sustain a new proto-spine, and as expected from the above analysis, this zone becomes narrower at higher bulk IRSp53. Overall, the tightest spine spacing was possible for smaller initiation delays and when there was enough IRSp53 to sustain them but not enough to get them to merge (Fig. 3 I). Existing proto-spines were more stable and restricted new spine formation.

### Glutamate triggered synaptic signaling pathways activate IRSp53

Next, we implemented a glutamate-triggered signaling pathway that is known to drive de-novo spine formation via calcium signaling, CaMKII, RhoGEF, and Rac-1, leading to modulation of IRSp53 activation (*IRSp*53_*act*_) (9, 35, 69– 71) (Fig. 4 A, Supplementary Fig.S2).

**Fig. 4.**
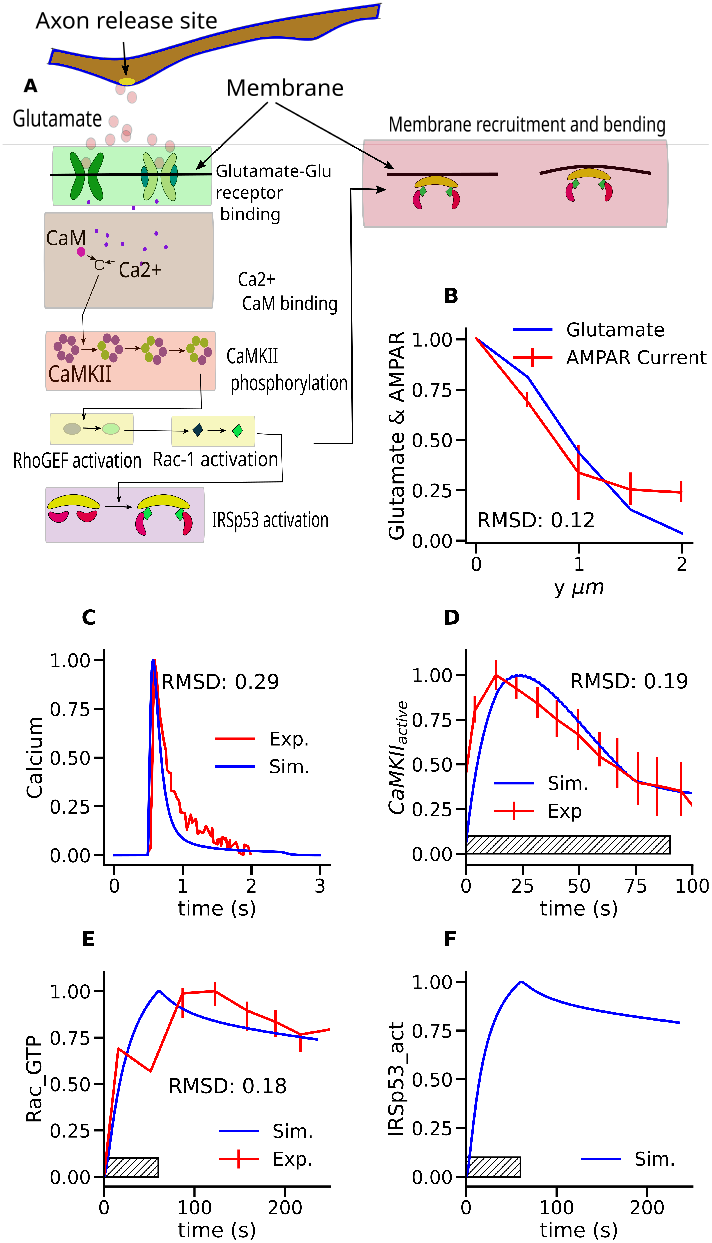
Signaling model match to experiment. Simulated (blue) and experimental (red) time courses of key molecules upon glutamate uncaging. All plots were normalized to their maxima. A) Model reaction block diagram B) Simulated glutamate concentration at dendrite as a function of the height *y* of release point above dendrite, compared to the experimental profile for AMPA receptor current as a function of release height (30). Pearson correlation value p = 1.06e-2. C) Experimental (72) and simulated Calcium time course measured postsynaptically following a single vesicle release of glutamate. p = 1.07e-5 D) Time course of CaMKII for 45 pulses of glutamate release at 0.5 Hz (30). p = 1.51e-3 E) Rac1 time course in response 30 pulses of glutamate at 0.5 Hz (73). p = 4.74e-4 F) Simulated time course of activated IRSp53, *IRSp*53_*act*_. The hatched block shows the duration of glutamate stimuli in experiments and simulations

We assembled a reaction-diffusion mass-action model using these pathways and used published experiments to systematically constrain rates for successive steps. We first used simple diffusion calculations (Eq. 17) to compare predicted glutamate concentrations as a function of distance from the dendrite (Fig. 4 B). This matched well with measurements of AMPAR current recorded postsynaptically at a range of distances from glutamate uncaging stimuli, where the current was used as a proxy for glutamate levels at the dendrite (RMSD = 0.12). We next compared model with experimental calcium dynamics (Fig. 4 C) (RMSD = 0.29), then CaMKII activation (RMSD = 0.19) (Fig. 4 D), and finally Rac-1 activation(RMSD = 0.18) (Fig. 4 E). All these model stages were good matches to experiment. We were not able to find direct readouts of IRSp53 as a function of glutamate input. Hence the final reaction step is assumed to be non rate-limiting, and predicts that the IRSp53 response time-course is similar to Rac1 responses (Fig. 4 F).

Combined, our model is a good fit to experimental observations of the space-time evolution of individual molecules in response to glutamate stimuli (14, 15, 29).

### Burst synaptic input gives stronger and broader IRSp53 activation than periodic input

We next used this model to compare IRSp53 buildup between burst and periodic stimuli, over space and time. Typical experiments on de-novo spine formation consist of glutamate pulses delivered at several frequencies, with the probability of spine formation increasing with frequency (9). We contrasted such periodic stimuli with burst input, as the latter is known to elicit selective synaptic responses (74). At all stages of the chemical pathway the burst (6 bursts, 10 pulses at 40 Hz per burst, inter-burst interval = 2s) input was much more effective than periodic stimuli of 60 pulses at 0.5 Hz (Supplementary Fig. S3). IRSp53 reached the threshold of 0.4 *µM* IRSp53 when the stimulus was within 3 *µm* of the dendrite, whereas the peak IRSp53 for periodic input reached only 0.125 *µM* (Supplementary Fig. S3).

### Proto-spine distribution depends on input synchronization and is sensitive to ASD mutations

We next coupled our mechanical and chemical models to study how the spatio-temporal organization of stimuli affects proto-spine stability. We first considered interactions between two proto-spines; one already existing, and the other initiated with a burst input after a 5-second delay, while we varied the spacing between the two. We took a snapshot of spine survival at t = 60 seconds. We found that rapid burst input (40 Hz) and a spacing *>* 0.75 *µm* were needed to sustain both spines, but if the spines were too close (*<* 1.0*µm*, inter-burst interval *<*= 0.5s), proto-spines merged (Fig. 5 A, supplementary movie S3 A, B). The survival of the second proto-spine reduced as the inter-burst interval increased. Eventually, both proto-spines vanished when the second proto-spine was initiated *>*= 12*µm* from the existing proto-spine, and bursts occurred at an inter-burst interval *>*= 2.0 s. We also found an intermediate inter-burst interval range where the second proto-spine stabilized at a distance = 14.0 *µm* (supplementary movie S4 B), while the existing one vanished. This occurred because the stimulus generated sufficient IRSp53 at the location of the second proto-spine to sustain it, but there was insufficient diffusive spread for the first proto-spine to continue.

**Fig. 5.**
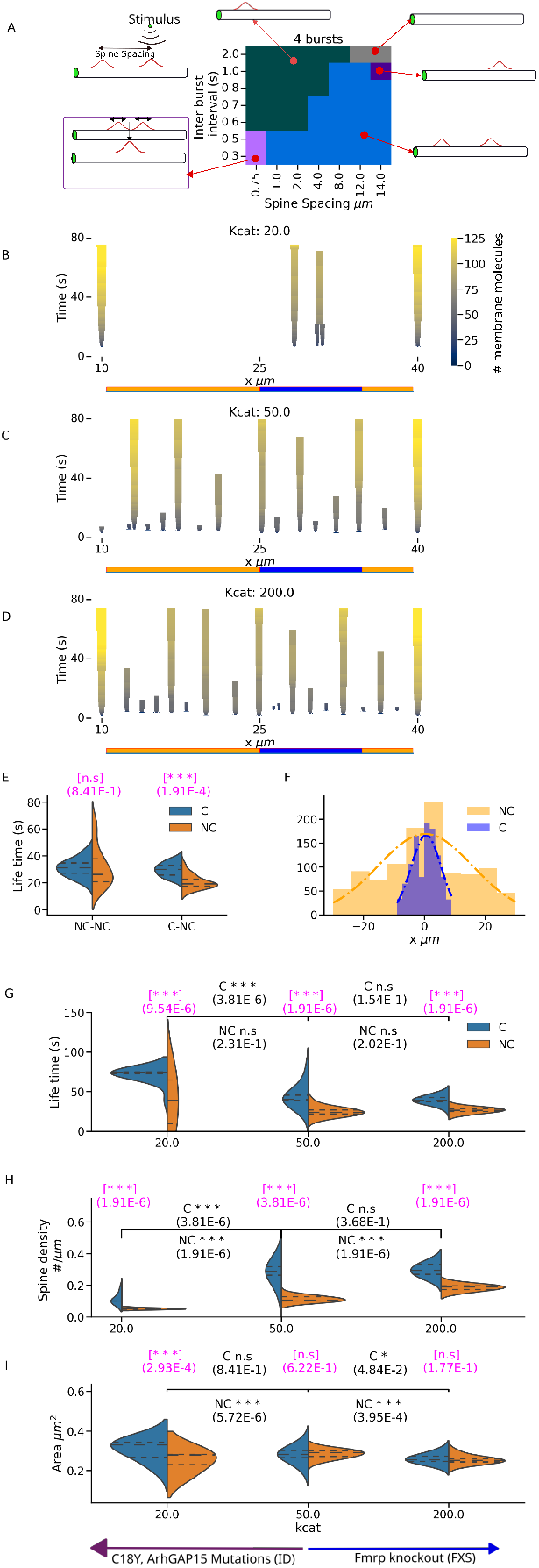
Dendritic spine distribution arises from mechano-chemical coupling and depends on signaling rates affected by ASD spectrum mutations. A) Multiple outcomes of proto-spine creation, removal, duplication or merging due to burst stimulus near existing proto-spine. A stimulus of four bursts was delivered at various inter-burst intervals and spine spacings from the existing proto-spine. B. C, and D) Spine life-cycle in the presence of ongoing non-correlated (orange) and correlated (blue), randomly positioned glutamate release near dendrite. The value of the kcat for the RhoGEF-catalysed Rac-1 activation was 20/s (B), 50/s (C, reference value) and 200/s (D). E) Control for effect of placement of correlated activity. Activity in both regions is a Poisson train with mean frequency of 1 Hz/*µm*. We compared lifetimes of proto-spines between condition 1 (NC-NC on the x axis), where non-correlated (NC) activity is delivered at both blue and orange regions, against C-NC, having correlated (C) activity in the blue region. There is no significant difference in the lifetimes of proto-spines in NC-NC, but in C-NC p=1.9e-4. F) Spatial correlation of inputs to C is much tighter than for NC. Distribution of distance (x) between two successive inputs for C and NC regions. KS Test p = 1.9e-32. G, H, and I show proto-spine lifetime (G), density (H), and area (I) across different Rho-GEF kcats, compared between C and NC regions of the dendrite. The arrows at the bottom show the direction of effects of possible ASD-related mutations which modulate the activity of Rac-1. p-values in pink are comparisons between C and NC for a given condition, and in black are comparisons between conditions.

It has been suggested that synchronized or highly correlated activity may lead to spine cluster formation (75). To explore this, we stimulated a 30 *µm* stretch of a dendrite along which 30 synapses were distributed uniformly. Within the dendrite, we embedded a 10 *µm* clustered region where synaptic input was more correlated (the C zone, blue). The C zone was bracketed by the zone of lower correlation (the NC zone, brown) of 15 *µm* and 5 *µm*. In each zone we delivered independent Poisson-timed bursts such that each successive burst arrived on a randomly chosen synapse. The mean firing rate for each region was 1 Hz/*µm*. Since zone C is more compact and has fewer inputs, its activity is more correlated than zone NC. We extracted the average lifetime, density, and size distribution for C and NC zones from the proto-spine distribution that emerged from mechano-chemical signaling downstream of this synaptic input.

We first asked if proto-spine lifetime distributions depended on input correlations (Fig. 5 E). The proto-spine lifetime was significantly longer in the C zone (Condition C-NC on x axis). As a control, we treated the entire dendrite as a single zone receiving Poisson burst input at 1 Hz/*µm* (Condition NC-NC on x axis). Here, lifetimes in the spatial regions assigned to the C and NC zones were similar.

Finally, we examined how proto-spine properties depended on signaling parameters sensitive to ASD-related mutations. Specifically, RhoGEF activity is a key readout of mutations associated with ASD (21, 76, 77). RhoGEF activates Rac-1, which is a converging point for various neurological disease-related pathways, and a key regulator of IRSp53 insertion in our model. Distinct mutations may raise or lower Rac1 activity, leading to bidirectional alterations in spine density(78, 79). The GEFD1 mutation in a RhoGEF called Trio causes reduction in RhoGEF-mediated activation of Rac-1 (80), whereas mutations in the seventh and eighth spectrin repeats cause its elevation (77, 80). Fragile X syndrome-related Fmrp mutations increase the activation of Rac-1 (81– 83), but conversely Rac-C18Y and ArhGAP15 mutations in intellectual disorders reduce Rac-1 activity (84, 85). To model these effects, we varied the catalytic rates for RhoGEF in catalyzing Rac-1 GTP formation (Supplementary Fig. S2D, Reaction number 6), between *kcat* = 20*/s*, to *kcat* = 200*/s* to bracket the experimentally fitted value of *kcat* = 50*/s* (Fig. 4). Correlated input (C) led to longer spine lifetimes than non-correlated (NC), over all values of *kcat*, and lifetimes were also longer at reduced *kcat*, e.g, GEFD1 mutations. (Fig. 5G).

Proto-spine density was always higher for correlated input (C) and it increased with *kcat* (Fig. 5 H, Supplementary Fig. S4). This parallels the phenotype of dense filopodia-like spines in the FXS variant of ASD, where FMR1 knock out leads to longer and thinner spines (86–91).

## Discussion

We have analyzed how membrane-binding proteins self-assemble into stable membrane deformations (proto-spines), with a rich repertoire of spacing and lifetimes depending on competition and input patterns. We first developed a mechanical-chemical model of stimulus-induced spine formation at the early stage of membrane protrusion before actin elongation or branching. Our mechanical model predicts stable proto-spines with biologically plausible spacing, spine competition, and merging of spines. These outcomes depend on IRSp53 protein levels, initiation timing, and spine position. We implemented a model of chemical signaling pathways to couple glutamate stimuli to IRSp53 and thence to the mechanical model. This showed that bursty and correlated input was most efficacious in triggering spine formation. Finally we merged the models to study spine stability, size, and spacing for different input patterns and for mutations linked to ASD.

### From coupled mechano-chemical signaling, to spines

Existing mechano-chemical models emphasize either the mechanical, or the chemical, but rarely integrate both. Many models focus on protein-membrane-lipid coupling, which gives rise to membrane deformation (92, 93), without exploring how the protein is recruited to the membrane, though this step has separately been modeled in terms of chemical potentials (94). A more detailed model of membrane deformation implemented a reaction-diffusion system involving N-WASP, Cdc42, and F-BAR driving F-BAR recruitment to the membrane (95). Membrane deformation was modeled as a height function resulting from F-BAR protein-membrane interaction energies and surface tension, explaining cortical membrane wave propagation in response to antigen stimulation of immune cells, but with an abstract modeling of chemical signaling and the mechanical feedback to it. Reaction-diffusion chemistry has also been used to understand the effect of different membrane shapes on the membrane protein distribution (96).

A critical maturation step in spine formation is the handover from proto-spine deformations to actin-induced spine stability, which involves proteins like Homer and Shank. There is a gap in the modeling literature for these handover steps. Rangamani et al. (28) have examined actin elongation and branching coupled to chemical signaling in the mature spine head without accounting for the mechanical energy rules that sculpt the spine. In membrane deformation studies (97, 98), mechanical energy was included, but the biochemical pathways were absent. Thus our study synthesizes and substantially extends several approaches to membrane deformation, by linking a novel formulation of proto-spine formation to its triggers from receptor signaling. This formulation yields numerous points of contact with experiment, as discussed below.

### Proto-spine dynamics replicate features of synapse elimination and turnover

Chen et al. (99) show that newly formed spines in the macaque prefrontal cortex pyramidal neurons are less stable than the mature preexisting spines. We extend this to the spine precursor domain through our finding that actin-less proto-spines also have a brief lifetime of about one minute in the absence of sustained input (Fig. 5). This duration of one minute is also consistent with the time-scale of de-novo spine formation (9). Chen et al also found that newly formed spines between two pre-existing spines will survive depending on the distance from the pre-existing spine: the larger the distance, the better the survivability (99). Consistent with this, our simulations show that proto-spines compete with pre-existing ones and are more likely to survive if further away (Fig 3 I, 5 A).

Synapse elimination is a further major mechanism for spine placement. For example, spine cluster formation has been found to be an outcome of in vivo-like synaptic activity (10). By applying a simple heuristic rule to delete spines that don’t respond to stimuli and inserting those that do, their model formed spine clusters. Synapse depression occurs where activity in a cluster is out of sync between neighboring synapses (100). Synapse elimination by competition has been known experimentally to result from proBDNF signaling (101), which is activity dependent. We obtain similar results for spine elimination and clustering from more physically-motivated calculations (Fig. 3, 5 D). Our simulations suggest that some of the competition dynamics for synapse elimination may occur even before the spines have matured.

### Activity-dependent proto-spine distributions in health and disease

Numerous experiments have characterized spine distributions as a function of cell type and location within the brain. Mature spine spacings can vary from 0.25 - 3 *µm* (7, 99). The spine spacing in pyramidal neurons in the prefrontal cortex of the Macaque is 2 - 4 *µm*, and in human temporal cortex is 2 *µm* (102). Our findings (Fig 3) show that competition among proto-spines establishes a proto-spine spacing of 1-3 *µm*, which is remarkably consistent with these experimental values despite the simplicity of our model. We speculate that the spacing templates laid down by proto-spines may persist through spine maturation.

Relatively few theoretical studies have examined factors which influence spine distribution. A study has shown that electrical potential is affected by the spine density, and spine density, in turn, is affected by the stem resistance because the decrease in stem resistance makes a higher number of spines electrically coupled to the dendrite (103). Turing patterning through reaction-diffusion patterning has been invoked to explain spine patterning (104), but this leads to uniform spine spacing, in contradiction to experimental observations (105). We find that wave-pinning models are better mapped to our mechano-chemical equations for spine initiation (Fig 2). These models possess more realistic properties of stimulus-driven spine placement and nonuniform spacing (Fig 2, 3, 5). This is consistent with observations to show that dendritic clustering of spines is input correlation dependent (75).

The activity-dependent placement of spines has major effects on subsequent cellular computation. Key readouts of learning are embedded in spine shape and distribution (10, 106, 107). Poirazi and Mel (106) have looked at the spine distribution and its effect on dendritic computation, but with a given equidistantly spaced spine distribution. Spine distribution has also been studied as a factor that affects the trafficking of AMPARs in the dendrite (108). Santamaria and De Schutter (109) have used spatial simulations to show that spine distributions affect the range of dendritic signaling, with implications for dendritic computation.

There are numerous examples of synaptic activity and signaling triggered changes in spine distributions and turnover. Our model provides mechanistic insights to bridge the steps between inputs and spine dynamics. We show that spine density and placement is elevated by correlated inputs (Fig. 5) and further that signaling changes in fragile X mutations lead to alterations in spine density (21, 88). While our work focuses on spine initiation, we suggest that these results have implications for phenomena such as spine turnover and density changes in long-term plasticity (110), as shown by the dependence of proto-spine lifetime on the degree of correlation of the stimulus (5 G and H). We suggest that known modulators of spine density such as stress (111–113), fear (114, 115) and enriched environment (12, 13) may provide signaling regulation of the pathways in our model through modification of activity of Rac-1 and RhoGEF, and thus lead to alterations in proto-spine spacing (Fig. 4A, 5 H). Overall, our analysis integrates a wide range of experimental and physiological correlates of spine dynamics and placement through an account of spine formation, competition, and stimulus-driven resource availability.

## Materials and Methods

### Mechanical deformation model

We employ an axisymmetric model of the proto-spine on the dendrite, with just three variables *R*^*d*^, *R*^*s*^ and *θ*, to describe the spine as a section of a sphere protruding from a locally flat dendrite (Fig. 1 A). The membrane shape has two curvatures of opposite sense: *R*^*d*^ is the radius of dome curvature (negative), and *R*^*s*^ is the radius of saddle curvature (positive). The area of the dome is *A*^*d*^ = 2 ∗ *π* × (*R*^*d*^)^2^ × (1 − *cos*(*θ*)) and the saddle area is *A*^*s*^ = 2 × *π* × *R*^*s*^ × (*r*_0_ × *θ* + *R*^*s*^ × (*cosθ* − 1)). The half length of the base of proto-spine is *r*_0_ = (*R*^*d*^ + *R*^*s*^) × *sin*(*θ*). The protein has a saturation density *ϕ*_*sat*_ = 1*/*50*nm*^2^. Surface fraction covered by proteins *ϕ* = *ϕ*_*tot*_/(*A*^*d*^ × *ϕ*_*sat*_).

We assume a separation of time-scales between mechanical relaxation of deformation (fast) and the exchange, diffusion and reactions in the model (slow). We perform energy minimization for the fast mechanical relaxation processes on every time-step of the numerical integration for the slower processes. The simplicity of our model makes this numerical approach computationally tractable, and allows us to calculate membrane shape evolution in response to dynamic chemical stimuli. Our model introduces chemical coupling to the mechanical system through a chemical potential *E*_*c*_ that modifies the membrane binding affinity of IRSp53. We also take into consideration the size requirement that the spine precursor has to reside on the dendrite of around 1 *µm* in diameter. A geometric constraint *M*_*c*_ penalizes unrealistically large spines. This term was negligible except for some of the shallow protrusions. In contrast to a recent report on IRSp53-induced membrane protrusion with a fixed size (48), our model allows for a wide range of possible sizes and shapes at the spine initiation stage.

### Energy terms

The total energy consists of bending energy of dome *E*_*BD*_, IBAR-dome curvature mismatch energy *E*_*m*_, Entropy *E*_*en*_, Aggregation energy *E*_*ag*_, saddle bending energy *E*_*SD*_, Membrane tension *E*_*t*_, the chemical energy due to difference in concentrations between dendrite and proto-spine *E*_*c*_, and the penalty term to prevent the proto-spine size from exceeding the dendritic radius *E*_*constraint*_ (Equations 1 - 10) *k*_*hat*_ and k are protein-modulated membrane rigidity (60 *k*_*B*_ *T*) and pure membrane rigidity (40 *k*_*B*_ *T*) (in N m) (116), respectively. *σ* = 5.5*µN/m* (117) is the membrane tension (in N/m).

The saddle bending energy has two components from two different curvatures *R*^*s*^ and *r*_*ψ*_ = *R*^*d*^ × *sin*(*θ*) (Fig. 1 A). These curvatures result in two bending energies: the Gaussian bending energy *E*_*sg*_, and the mean curvature energy *E*_*sh*_.

*µ*_0_ is the basal membrane affinity for IRSp53 (58), [*M*] and [*C*] are the membrane and dendritic concentrations. *m*_*t*_ = 5*nm* is the membrane thickness. *µ*_*mech*_ is the rate of change of the energy with respect to the change in *ϕ*_*tot*_ (Supplementary Eq. S1 - S4)

### Shape optimization

We expressed energy terms (Equations 1 - 10) as functions of the surface fraction of IRSp53 *ϕ*, which, in turn, is a function of the total number of IRSp53 in the membrane (*ϕ*_*tot*_) and geometric variables (Fig. 1 A). Thus,

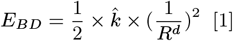

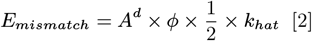

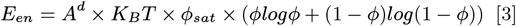

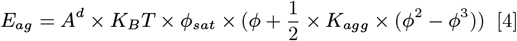

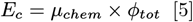

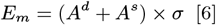

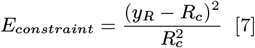

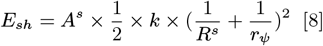

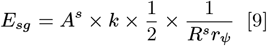

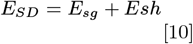

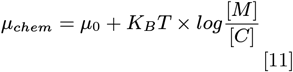

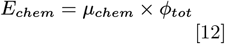

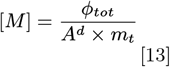

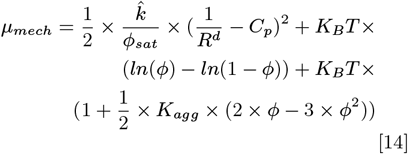

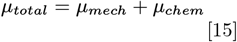

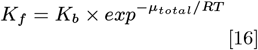

the total energy is a function of shape and the membrane rapidly converges to the shape with minimal total energy. To initiate a spine, we create a dummy protrusion of 0.5 *µm* width, i.e., *r*_0_ with just one molecule corresponding to *ϕ*_*tot*_ = 1. Following this, energy minimization governs further insertion of IRSp53 and membrane deformation. IRSp53 is recruited to the membrane due to its basal chemical affinity (−4.5 kcal/mol), thus increasing *ϕ*_*tot*_ in the dome region as per Equation 16. The shape is updated at each 200 ms interval by minimizing energy with respect to the *ϕ*_*tot*_, i.e, finding the set of geometric parameters corresponding to the minimum of the total of all energy terms (Eq. 1 - 12). Eq. 16 governs the recruitment of activated IRSp53 to the membrane. At each time-step, the distribution of membrane IRSp53 is programmatically made uniform along the dome region to match the assumption that *ϕ* is uniform inside the dome. The new shape modifies the chemical affinity of IRSp53 through changes in the membrane concentration [*M*] (Equation 11), and this is used for subsequent stages of the numerical integration.

### Simulation

We modeled the dendrite as a uniform cylinder, divided into 20 nm chemical compartments, coupled by diffusion. On each compartment we incorporated either the IRSp53 molecule only (Figures 1-3), or IRSp53 plus the entire chemical system (Figure 5). We used the multi-scale object-oriented simulation environment (MOOSE) to perform reaction-diffusion calculations (118). We created a simple ball-and-stick model for soma and dendrite, including NMDAR, AMPAR, and calcium channels on the dendrite. We modeled glutamate binding to the receptors and subsequent calcium influx (supplementary material Eq. S9-20). The incoming calcium triggers the chemical pathway downstream, activating IRSp53 (Supplementary Fig. S3). The activated IRSp53 is the input to the mechanical model.

In all simulations, spine initiation involves creating a dummy spine with no molecules (*ϕ*_*tot*_ = 0) and then allowing it to evolve according to the potential (Eq. 15 - 16).

In simulations with multiple proto-spines, the spines were initiated 200 ms apart (Fig. 3 A and Fig. 5 A) to prevent numerical artifacts that could arise from sharp concentration gradients.

### Poisson stimuli

In both C and NC regions, stimulus timings were chosen from a Poisson distribution, and every time a stimulus event occurred, it was placed randomly (uniformly) at any of the release sites along the region. The spacing between release sites was 1 *µm* for both regions. The NC region was twice the spatial extent of the region C. Thus, to ensure that each of C and NC received the same frequency of stimulus per unit length (1 Hz/*µm*), the mean Poisson frequency for C was half that for NC in panels B to I in Fig. 5. As a corollary of the smaller length of C, the spatial correlation of its input was much tighter than for NC (Fig. 5 F).

### Glutamate molecule distribution

The distribution of gluta-mate (*c*(*x, y, t*)) in the 2D space between the stimulus source and the dendrite is modeled as Eq. 17 (119), where *D*_*glu*_ is the glutamate diffusion constant, h is the distance between the stimulus source and the dendrite, and *N*_*glu*_ is the number of glutamate molecules at the stimulus source for a single pulse. We set *N*_*glu*_ to 8000, corresponding to the number of glutamate molecules in a single vesicle (119), and *D*_*glu*_ to 300 *µm*^2^*/s* (120).

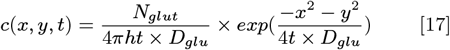

### Turing and Wave-pinning model

The Turing model used in Fig. 2 A was obtained from (121). The wave-pinning model was from (122). Both the Turing and wave-pinning models used reaction-diffusion equations for spatio-temporal evolution of two species A and B, in which molecule A was assumed membrane bound with diffusion coefficient 0.1 *µm*^2^*/s* and the cytosolic molecule B diffused more rapidly at 10 *µm*^2^*/s*. The stimulus in both cases was a Gaussian distribution of molecules, corresponding to a release of 50 molecules from a source at height 0.1 *µm* from the 1-dimensional reaction system. The diffusion coefficient of the stimulus was set to 10 *µm*^2^*/s*. This stimulus waveform (*A*_*stim*_) was added to the evolution equation for molecule A for 2 seconds as the source of perturbation (Eq. 18 for Turing pattern and Eq. 19 for wave-pinning) in Fig. 2 A.

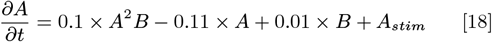

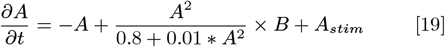

In Fig. 2 B, the increase in the stimulus intensity from top to bottom was implemented as 1, 2, and 3 times the stimulus intensity used for 2 A.

## ACKNOWLEDGMENTS

We thank Drs. Deepak Nair, Pramod Pullarkat and P. B. Sunil Kumar for useful discussions. VVP was supported by SERB N PDF PDF/2017/000419 and India Alliance grant IA/TSG/22/1/600405 and a Institute postdoc from NCBS. This work was carried out at NCBS-TIFR which is supported by the Department of Atomic Energy, Government of India, under Project Identification No. RTI 4006.

