## Supplementary material for "Formation, stability, and competition in dendritic proto-spines": Supplimentary material

**This PDF file includes:**

Supporting text
Figs. S1 to S4
SI References

### Supporting Information Text

In this material, we show the effect of model parameters on the energy of proto-spines, the details of chemical kinetics used to generate Figs. 4 and 5 in the main text, and the investigation of spatio-temporal evolution of key molecules in the glutamate-IRSp53 pathway.

### Parameter sweep for aggregation coefficients

In this section, we explore the parameter space from which we selected the values of model parameters used in the main text. The mechanical energy of the membrane protrusion has two unknown parameters: $k_{agg}$, which determines the aggregation strength of IRSp53, and $\mu_0$, which determines the basal binding affinity of IRSp53 to the membrane. $\mu_0$ is known to be in the range of -4 to -7 kcal/mol (1). The values we chose for the unknown parameters are shown in S1 A. We chose the threshold concentration of IRSp53 to be 0.4 $\mu M$ because the difference between the energy from wave speed analysis (Supplementary methods) and the absolute energy minimum $E_{sharp}$ (S1 B) is negligible beyond that concentration for all aggregation coefficients (Fig. S1 D). The threshold of 0.4 $\mu M$ IRSp53 to trigger spine initiation also is consistent with Fig. 2H, based on the stipulation that proto-spines should have a diameter of at least 0.5 $\mu m$, and the reference aggregation rate of -60 $K_B T$. Using this constraint for threshold concentration as a function of aggregation energy, we found that the theoretical calculations matched the simulation (Fig. S1 D).

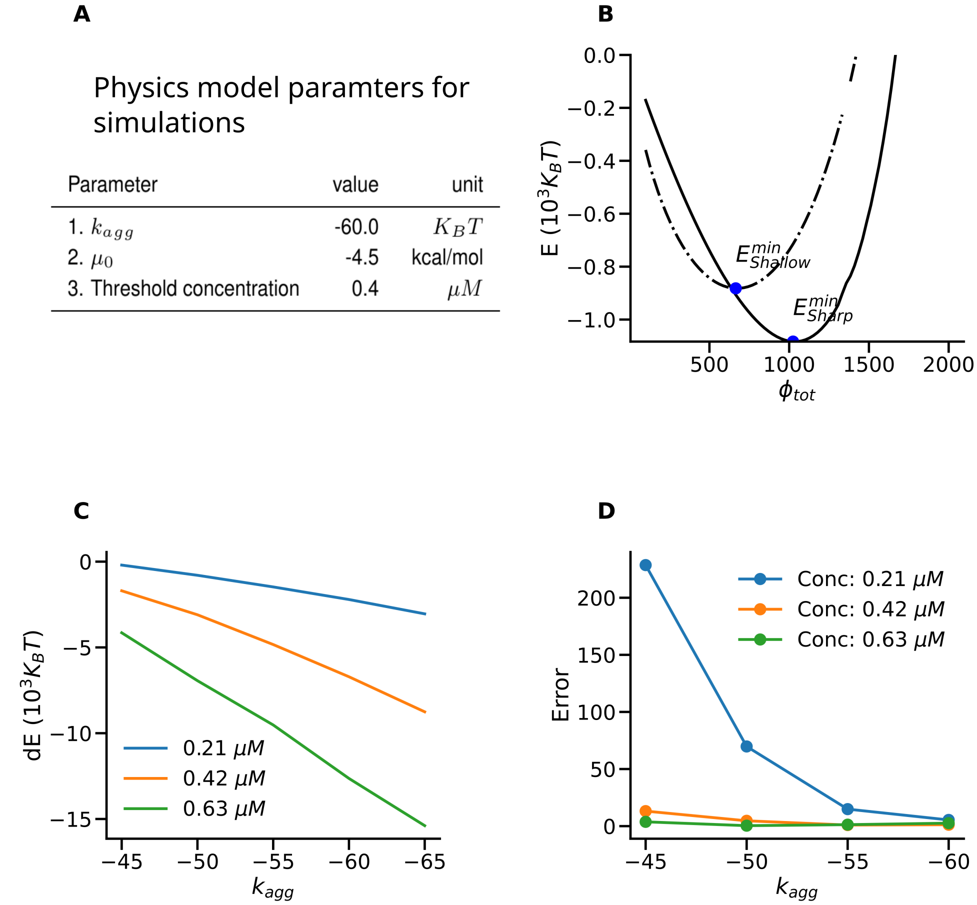


| Parameter | value | unit |
|---|---|---|
| 1. $k_{agg}$ | -60.0 | $K_B T$ |
| 2. $\mu_0$ | -4.5 | kcal/mol |
| 3. Threshold concentration | 0.4 | $\mu M$ |

**Fig. S1.** Parameter sweep across aggregation coefficients and concentrations. A) Parameters used in simulations and wave-speed analysis. B) The energy branches for shallow and sharp protrusions. C) The variation of $dE$ with respect to the aggregation coefficients. $dE = E^{min}_{sharp} - E^{min}_{shallow}$. D) The difference between the optimal energy from wave-speed analysis and the absolute energy minimum $E_{sharp}$. $Error = abs((E^{analysis}_{wave} - E_{sim})/E^{analysis}_{wave}) \times 100$

**A**

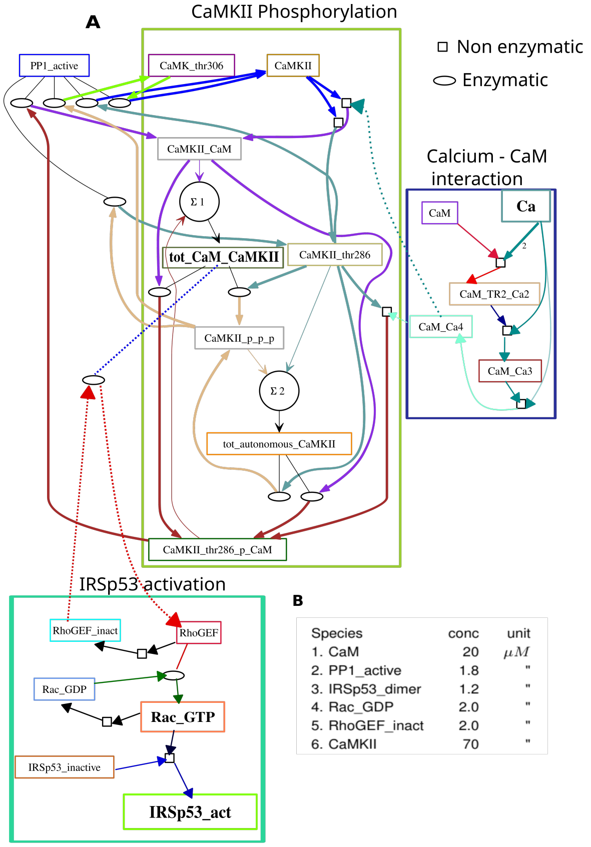


**B**

| Species | conc | unit |
|---|---|---|
| 1. CaM | 20 | $\mu M$ |
| 2. PP1_active | 1.8 | " |
| 3. IRSp53_dimer | 1.2 | " |
| 4. Rac_GDP | 2.0 | " |
| 5. RhoGEF_inact | 2.0 | " |
| 6. CaMKII | 70 | " |

**C**

Non-enzymatic reactions

| Reactions | Kf | unit | Kb | unit |
|---|---|---|---|---|
| 1. CaM + 2 Ca <===> CaM_TR2_Ca2 | 71.99 | $1/(\mu M^2 * s)$ | 72 | 1/s |
| 2. CaM_TR2_Ca2 + Ca <===> CaM_Ca3 | 3.6 | $1/(\mu M * s)$ | 10 | 1/s |
| 3. CaM_Ca3 + Ca <===> CaM_Ca4 | 0.47 | $1/(\mu M * s)$ | 10 | 1/s |
| 4. CaM_Ca4 + CaMKII <===> CaMKII_CaM | 50 | $1/(\mu M * s)$ | 0.1 | 1/s |
| 5. CaM_Ca4 + CaMKII_thr286 <===> CaMKII_thr286_p_CaM | 1000 | $1/(\mu M * s)$ | 0.1 | 1/s |
| 6. RhoGEF <===> RhoGEF_inact | 0.2 | 1/s | 4 | 1/s |
| 7. Rac_GTP <===> Rac_GDP | 0.2 | 1/s | 0.2 | 1/s |
| 8. Rac_GTP + IRSp53_dimer <===> IRSp53_active | 2 | $1/(\mu M * s)$ | 4 | 1/s |

**D**

Enzymatic reactions

| Reactions | $K_m$ | unit | $k_{cat}$ | unit |
|---|---|---|---|---|
| 1. CaMKII_thr286_p_CaM —PP1_active–> CaMKII_CaM | 0.55 | $\mu M$ | 0.35 | 1/s |
| 2. CaMKII_p_p_p —PP1_active–> CaMKII_thr286 | 5.55 | " | 0.35 | " |
| 3. CaMK_thr306 —PP1_active–> CaMKII | 5.55 | " | 0.35 | " |
| 4. CaMKII_p_p_p —PP1_active–> CaMK_thr306 | 5.55 | " | 0.35 | " |
| 5. PP1_active–> CaMKII | 5.55 | " | 0.35 | " |
| 6. Rac_GDP —RhoGEF–> Rac_GTP | 80.6 | " | 50 | " |
| 7. CaMKII_thr286 —tot_CaM_CaMKII–> CaMKII_p_p_p | 123.65 | " | 6 | " |
| 8. CaMKII_CaM —tot_CaM_CaMKII–> CaMKII_thr286_p_CaM | 123.65 | " | 0.5 | " |
| 9. RhoGEF_inact —tot_CaM_CaMKII–> RhoGEF | 50.37 | " | 4 | " |
| 10. CaMKII_thr286 —tot_autonomous_CaMKII–> CaMKII_p_p_p | 190.47 | " | 0.5 | " |

**Fig. S2.** The biochemical pathway activating IRSp53 and their kinetics parameters. A) The detailed pathway starting from the calcium entry to the IRSp53 activation. The pathway is organised into three groups: 1) The calcium (Ca) entry and Calcium-Calmodulin (CaM) binding. 2) CaMKII phosphorylation showing the steps of Calcium-Calmodulin dependent phosphorylation of Calcium-Calmodulin Kinase 2 (CaMKII), which then acts as the activator of RhoGEF, 3) IRSp53 activation group modulated by RhoGEF and Rac-1. B) Initial concentration of molecules in the pathway. C) and D) are the chemical kinetics parameters of the non-enzymatic and enzymatic reactions, respectively.

 

### Spatio-temporal evolution of key molecules in response to a stimulus

The efficiency of chemical signaling in terms of activating key molecules varies according to the type of stimulus. Fig. S3 shows the response of the same biochemical pathway (Fig. S2 A) to two different stimuli, i.e., a sparse stimulus of 60 single pulses and a concentrated stimulus of 6 bursts, with each burst consisting of 10 pulses. The activated IRSp53 is five times higher for the burst stimulus when compared to the single pulses. The calcium entry is also higher for the burst stimulus.

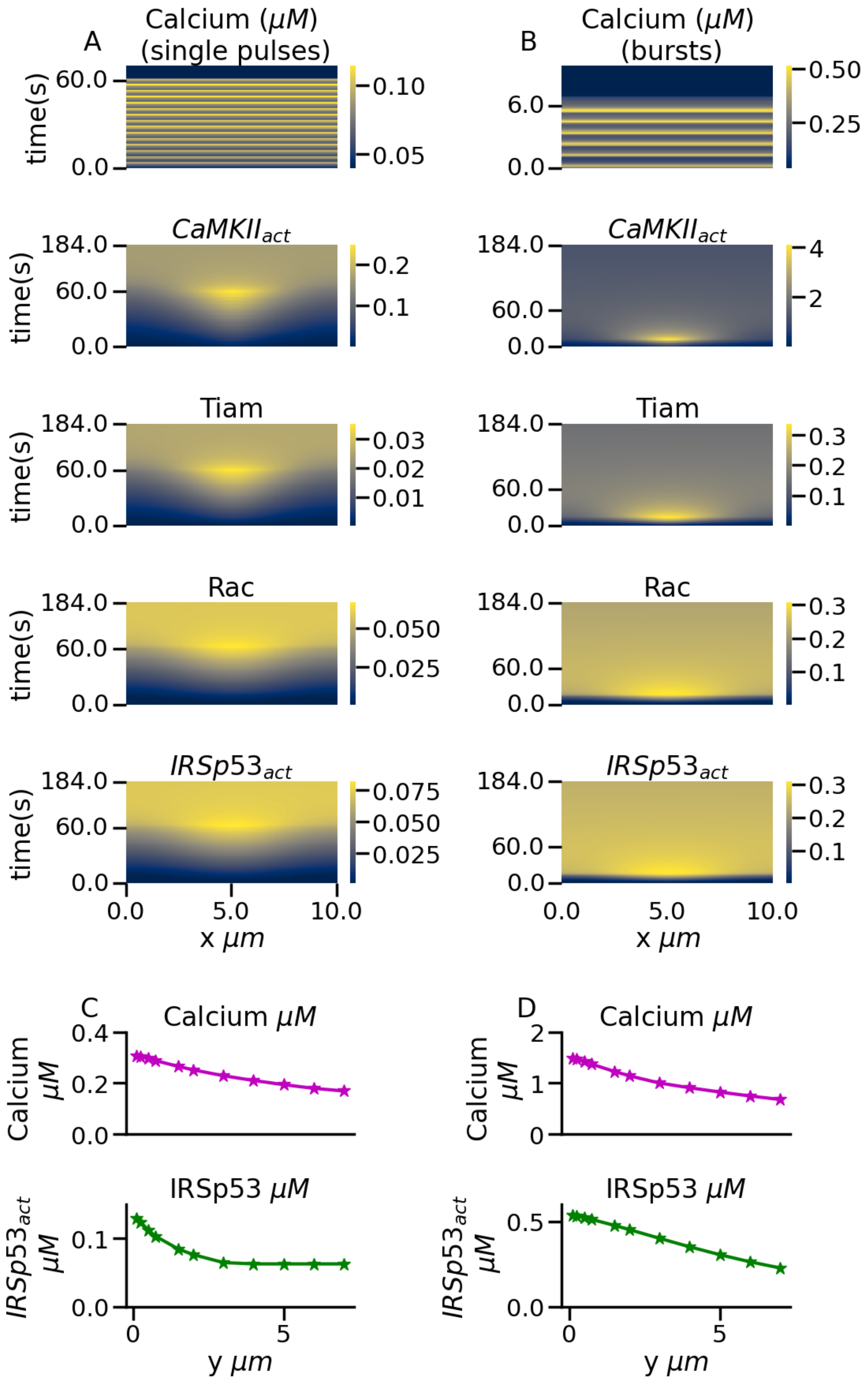


**Fig. S3.** Performance of biochemical pathway for single pulses Vs bursts. A) and B) columns are for single pulses and bursts, respectively. The same number of pulses for each column, i.e., 60 pulses at 1 Hz for a single pulse case and six bursts separated by 1 s, each containing 10 pulses at 40 Hz for a burst case. C) and D) show the activation of Calcium and IRSp53 for single pulses and bursts, respectively.

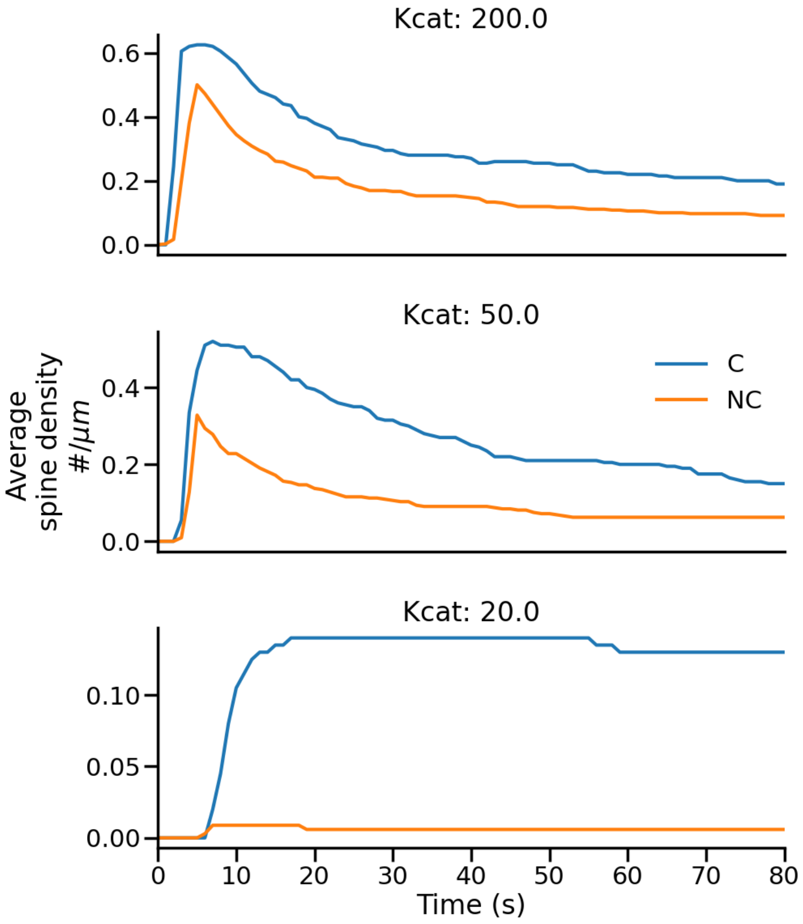


**Fig. S4.** Time evolution of spine density averaged over 20 trials for F = 10 Hz and various RhoGEF kcats

**Methods.** The mechanical bending and protein-membrane interaction in the proto-spine induces a mechanical potential ($\mu_{mech}$) that acts along with the chemical potential $\mu_{chem}$ to recruit the molecules to the membrane.

$$E_{dome} = E_{mismatch} + E_{en} + E_{agg} + E_c \quad [1]$$

$$\mu_{mech} = \frac{dE_{dome}}{d\phi_{tot}} = \frac{dE}{d\phi} \times \frac{d\phi}{d\phi_{tot}} \quad [2]$$

$$\mu_{mech} = \frac{1}{2} \times \frac{\hat{k}}{\phi_{sat}} \times (\frac{1}{R^d} - C_p)^2 + K_B T \times (ln(\phi) - ln(1-\phi)) + \quad [3]$$

$$K_B T \times (1 + \frac{1}{2} \times K_{agg} \times (2 \times \phi - 3 \times \phi^2)) \quad [4]$$

$$\mu_{total} = \mu_{mech} + \mu_{chem} \quad [5]$$

Wave-pinning models describe a pattern of peaks and valleys, with peaks representing the presence of molecules, like the Turing patterns. The stability of wave-pinning models is formulated as a function of the wave speed, i.e., the speed at which the size of the peak expands or shrinks (2). In our model, the proto-spine size changes to accommodate molecules in the dome. Therefore, we describe proto-spine size as a function of $\phi_{tot}$ (Eq. 6). Where $L = R^d \times sin(\theta)$ is the half length of the dome.

$$W_s = \frac{dL}{dt} = \frac{dL}{d\phi_{tot}} \times \frac{d\phi_{tot}}{dt} \quad [6]$$

The rate at which $\phi_{tot}$ changes depends on the rate at which IRSp53 is recruited to the membrane ($K_f$ and $K_b$), which, in turn, depends on the potential $\mu_{tot}$. We can then express the wave speed as Eq. 7.

$$W_s = (\frac{1}{2} \times \frac{1}{\sqrt{(\phi_{tot})}} \times C_I \times \sqrt{(1 + cos(\theta))} + C_I^2 \times \frac{cos(\theta)}{L}) \times (K_f[C] - Kb[M]) \quad [7]$$

$$C_I = \sqrt{(\frac{1}{2\pi\phi\phi_{sat}})} \quad [8]$$

As shown in Fig. 2E of the main article, the wave speed changes with $\phi_{tot}$. The energy corresponding to $W_s = 0$ is the optimal energy.

The calcium entry to the dendrite in response to the glutamate binding is modeled as per the equations below (3, 4). In these equations, $g$ is the conductance of channels with subscripts $gluR$ denoting AMPAR and $NMDAR$ NMDAR.

Eq. 20 describes calcium through the receptors.

$$g_{Ca} = g_{max} Cas^2 r \qquad [9]$$

$$g_{max} = 40.0 s/m^2$$

$$s - gate : \alpha = \frac{1.6}{1 + exp(-0.072 * (V - 65))} \qquad [10]$$

$$\beta = \frac{0.02(V - 51.1)}{exp(\frac{V-51.1}{5}) - 1} \qquad [11]$$

$$r - gate : V <= 0; \alpha = 0.005 \qquad [12]$$

$$\beta = 0.0 \qquad [13]$$

$$r - gate : V > 0 : \alpha = \frac{exp(-V/20)}{200} \qquad [14]$$

$$\beta = 0.005 - \alpha \qquad [15]$$

$$E_{Ca} = 140$$

$$g_{gluR} = \frac{A.g_{maxGluR}}{\tau_1 - \tau_2} \times (exp(\frac{-t}{\tau_1}) - exp(\frac{-t}{\tau_2})) \qquad [16]$$

$$g_{maxGluR} = 4000 s/m^2$$

$$tau_1 = 2, \tau_2 = 9$$

$$g_{NMDAR} = \frac{g_{maxNMDAR}}{\tau} \times exp(\frac{-t}{\tau}) \times \frac{K_{Mg}}{K_{Mg} + [Mg]} \qquad [17]$$

$$g_{maxNMDAR} = 800 s/m^2$$

$$K_{Mg} = exp(\frac{(V - E_{rest}) \times \gamma}{\eta}) \qquad [18]$$

$$E_{rest} = -60 mV$$

$$\tau = 20, \gamma = 0.28$$

$$I_{NMDAR_{Ca}} = g_{NMDAR}.Ca.ln(\frac{[Ca_{out}]}{[C_{in}]}).V.\frac{[Ca_{in}] - \phi[Ca_{out}]}{(1 - \phi)([Ca_{in}] - [Ca_{out}])} \qquad [19]$$

$$\phi = exp(\frac{-VFz}{RT}), F = 96485 sA/mol, z = 2, T = 300K, [Ca_{out}] = 1.5 mM, [Ca_{in}] = 0.08 \mu M$$

$$R = 8.314 J/Kmol$$

$$\frac{d[Ca]}{dt} = \phi(I_{Ca} + Ca_{NMDA}) - [Ca]/1.33 \qquad [20]$$
